# Salivary flow rate and prevalence of subjective halitosis in major depressive disorder patients

**DOI:** 10.1101/2025.04.26.650815

**Authors:** Ameer Ali Althabhawee, Taghreed Fadhil Zaidan

## Abstract

The global incidence of major depressive disorder (MDD) is steadily increasing, leading to a concomitant surge in the need for oral medication in developed nations. The field of oral medicine serves as an intermediary between the disciplines of medicine and dentistry, with a focus on the study and treatment of various disorders and orofacial pain. Previous research has established a connection between these conditions and psychiatric disorders. The study sample comprised 49 individuals who had been diagnosed with major depressive disorder and had been receiving treatment for a minimum of two weeks. The control group consisted of 34 healthy participants who exhibited no indications or symptoms of systemic disease. The patients received diagnoses in accordance with the Diagnostic and Statistical Manual of Mental Disorders, Fifth Edition (DSM-5). Reporting of subjective halitosis was highly significant (p< 0.000) in MDD patients than in control subjects, as for unstimulated salivary flow rate the mean of salivary flow rate in MDD patients was 0.39 ml/mints, with range of 0.03-1 ml/min, while for control subjects was 0.64 ml/mints with range of 0.03-0.6 ml/mint.

The results of this study showed that, there was a highly significant difference of salivary flow rate between control and study groups (p value <0.05 at p=0.000)

In conclusion, we found that major depressive disorder patients have low unstimulated salivary flow rate and the subjective halitosis is more prevalent in this group than in control group.

## 1. Introduction

Major depressive disorder (MDD), considered the leading cause of disability globally and presents with diverse manifestations, impacts approximately 20% of individuals [1]. A correlation has been observed between the symptoms of Major Depressive Disorder (MDD) and structural and neurochemical impairments in the corticolimbic areas of the brain [2]. The clinical presentation of depression in individuals residing in Baghdad exhibits similarities to that observed in individuals from various global regions [3].

Multiple studies have indicated that Iraq exhibits a significant prevalence of depression [4,5,6].

A recent study conducted in Iraq has identified depression as a significant criterion that had been previously disregarded. The study proposed that the inclusion of depression as a criterion may have the potential to facilitate the early identification of individuals with Behcet’s disease [7].

Halitosis, also known as oral malodor, is primarily attributed to the presence of volatile sulphur compounds that are generated by microorganisms residing in the oral cavity. The global prevalence of this condition ranges from approximately 22% to 50% [8].

More than 32 % of people have bad breath, according to a recent systematic review of the global prevalence [8] of this condition. The aetiology of malodour can stem from either systemic or local factors [8]. In the majority of instances, poor oral hygiene practises are strongly associated with self-reported bad breath [9].

According to the studies, psychological factors can also play a significant role in the prevalence of halitosis [10]. This association has been addressed with two distinct explanations. The use of medication to treat these conditions may alter the flow and composition of saliva, leading to an increase in bacterial colonisation and the degradation of proteins in the mouth, resulting in an increase in volatile sulfur-containing compounds responsible for bad breath [10,11]. In addition, people with mood disorders, such as those who are depressed, anxious, or stressed, may engage in less self-care, less motivated to maintain oral health, these individuals exhibit poor hygiene. [12].

Saliva is a mix of secretions from the major and minor salivary glands[13], which functions as an oral cleanser, aiding in the chewing of food and facilitating swallowing[14]. The buffering effect of saliva results in the neutralization of oral acids and the protection of the dentition. Furthermore, it is worth noting that saliva serves to strengthen the mucosal barrier and exhibits antimicrobial properties [15].

The unstimulated salivary flow rate refers to the quantity of saliva produced by the major and minor salivary glands within a one-minute period, in the absence of any external stimulation, the typical range of salivary flow rates in unstimulated and stimulated conditions is 0.3-0.5 and 0.5-0.7 mL/min, respectively. [14,16]. Lower values are indicative of impaired functioning of the salivary glands.

Reduced salivation can result in speech difficulties, chewing disorders, mucosal inflammation (mucositis), oral Candida infections, and mucosal atrophy. It can increase the buildup of plaque and reduce the buffering capacity of saliva [13].

## 2. Martials and Methods

This cross-sectional study was conducted at Al-Hakim Hospital, located in Najaf City, Iraq. Ethical approval for the study was obtained from the Ethical Committee of the College of Dentistry at Baghdad University, with the project being assigned the identification number 458722.

### 2.1 Data collection

The study comprised a study group of 49 patients diagnosed with Major Depressive Disorder (MDD), all of whom had been receiving treatment for a minimum duration of two weeks.

The control group comprises 34 individuals who are in good health and exhibit no indications or symptoms of systemic illness.

The study group was diagnosed using the Fifth Edition of The Diagnostic and Statistical Manual of Mental Disorders (DSM5) by psychiatric specialists at Al-Hakim Hospital in the city of Najaf.

### 2.2 Inclusion& exclusion criteria

In order to be eligible for participation in this study, individuals aged 18 years or older who have received a diagnosis of depression from a qualified psychiatrist are considered. The exclusion criteria encompassed individuals who sought medical attention for emergencies, were incapable of independently completing the questionnaire, were pregnant, were undergoing corticosteroid treatment, or had a history of radio or chemotherapy. All patients underwent a comprehensive oral examination to identify potential oral manifestations and were also queried regarding the presence of halitosis. The examined sample collection spanned from January 30, 2021, to April 29, 2022.

### 2.3 Salivary flow rate

The whole resting (unstimulated) saliva was collected from each subject by spitting method in dried, sterilized graduated plastic tube for 10 minutes; then the volume of saliva measured and placed in a test tube. The rate was measured as the “volume of saliva in (ml) divided by the time required for collection in (minutes)” [17].

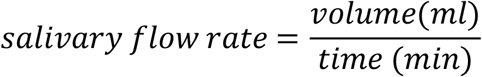

## 3. Results and discussion

The study revealed that individuals diagnosed with Major Depressive Disorder (MDD) exhibited a wide age range ranging from 23 to 66 years, whereas the control subjects, who did not have MDD, ranged in age from 20 to 57 years. The average age of individuals diagnosed with Major Depressive Disorder (MDD) was found to be 44.3 years, with a standard deviation of ±10.19 years. In the control group, the average age was found to be 41.26 years, with a standard deviation of ±10.98 years. There was no statistically significant difference observed in this regard. The study revealed that within the group of patients diagnosed with Major Depressive Disorder (MDD), there were 26 males (53.1%) and 23 females (46.9%). In comparison, the control group consisting of healthy individuals exhibited 19 males (55.9%) and 15 females (44.1%), Table (1).

**Table 1.** Frequency and percentage with halitosis in MDD patients and control subjects.

| Halitosis | Study Group MDD |  | Control Group |  | p-value |
| --- | --- | --- | --- | --- | --- |
|  | No. | % | No. | % |  |
| No | 16 | 30.6 | 32 | 94.1 | **0.000**<br>**\*\* (Hs)** |
| Yes | 33 | 67.3 | 2 | 5.9 |  |
| **Total** | **49** | **100** | **34** | **100** |  |

\*\*Hs: High significant

### 3.1 Halitosis

The results showed that for the study group 16 (30.6%) patients answered with no when asked about mouth odour, while 33 (67.3%) patients responded with yes.

As for control group 32 (94.1 %) individuals didn’t report mouth odour, while 2 (5.9%) individuals reported mouth odour. The results of this study showed that there was a highly significant difference (p≤ 0.001) of subjective halitosis between control and study groups. Table (1)

Genuine halitosis has been defined as halitosis confirmed by organoleptic tests and objectively assessed, whereas pseudo-halitosis or halitophobia can develop in cases where bad odour is not detected by this test as well as by other individuals.[18]

Although the organoleptic test has been considered the “gold standard” on halitosis evaluation, the use of a self-reported questionnaire has been used in the literature [10,11,12,19].

A recent meta-analysis comparing the prevalence acquired with clinical and self-reported measurements found that the method used to evaluate halitosis had no effect on the heterogeneity between studies [8].

The results of this study are in agreement with a recent study on this subject [20].

To establish a causal relationship between subjective halitosis and Major depressive disorder is beyond the scope of this study, however it could be suggested that two hypotheses can be assumed, firstly the perception of oneself in MDD patient is usually negative, and hence the reporting of halitosis, secondly, the bad oral hygiene practices for the patients of depressive disorder can lead to halitosis, further researches are needed in this area.

### 3.2 Unstimulated salivary flow rate

The mean of salivary flow rate in MDD patients was 0.39 ml/mints, with range of 0.03-1 ml/min, while for control subjects was 0.64 ml/mints with range of 0.03-0.6 ml/mint.

The results of this study showed that there was a highly significant difference of salivary flow rate between control and study groups (p value <0.05 at p=0.000), demonstrating significantly decreased salivary flow rate levels in MDD patients compared to the control group, using t-test table (2)

**Table 2.** The mean, standard deviation and range of salivary flow rate in MDD and control subjects.

| Groups | No. | Mean<br>mL/min | SDD | Range | p-value |
| --- | --- | --- | --- | --- | --- |
| patients | 49 | 0.39 | 0.24 | 0.03-1 | **\*\*0.000**<br>**(Hs)** |
| control | 34 | 0.64 | 0.29 | 0.03-0.6 |  |

\*\*Hs: high significant $p \leq 0.001$

The highest percentage of MDD patients (18.4%) were with saliva flow rate of 0.40, and of 0.60 ml/minuets, as shown in table (3)

**Table 3.** Frequency and percentage of MDD and control subjects according to salivary flow rate.

| SFR\Mint | Study Group |  | SFR\Mint | Control Group |  |
| --- | --- | --- | --- | --- | --- |
|  | No. | % |  | No. | % |
| 0.03 | 1 | 12.2 | 0.30 | 1 | 2.9 |
| 0.06 | 2 | 4.08 | 0.40 | 7 | 20.6 |
| 0.08 | 3 | 6.12 | 0.50 | 5 | 14.7 |
| 0.10 | 1 | 2 | 0.60 | 7 | 20.6 |
| 0.20 | 8 | 16.3 | 0.70 | 9 | 26.5 |
| 0.30 | 5 | 10.2 | 0.72 | 1 | 2.9 |
| 0.40 | 9 | 18.4 | 0.90 | 1 | 2.9 |
| 0.50 | 5 | 10.2 | 1.20 | 2 | 5.9 |
| 0.60 | 9 | 18.4 | 1.90 | 1 | 2.9 |
| 0.70 | 2 | 4.1 |  |  |  |
| 0.80 | 3 | 6.1 |  |  |  |
| 1 | 1 | 2 |  |  |  |
| **Total** | **49** | **100** |  | **34** | **100** |

Our study demonstrated that the salivary flow rate in the study group is significantly lower than that in control group consistent with the results of most previous studies in this area [21,22]

## 5. Conclusion

Major depressive disorder plays a substantial role in the reduction of the rate of salivary flow. This decrease in saliva flow rate can lead to multiple oral health problems in already distressed subjects. we recommend using artificial saliva, adjusting/changing antidepressant medication, if possible, furthermore depressed patients seemed to report more of subjective halitosis, the anti-depressant medications also cause a significant decrease in salivary flow rate, decreased salivary flow rate leads to multiple oral health problems in an already diseased patients, so, it is recommended to use an artificial saliva, and adjusting/ changing antidepressant medication if possible. Furthermore, depressed patients seem to report subjective halitosis.

